# NT-C2–Dependent Phosphoinositide Binding Controls PLASTID MOVEMENT IMPAIRED1 Localization and Function

**DOI:** 10.64898/2025.12.30.697064

**Authors:** Dominika Cieślak, Zuzanna Staszałek, Paweł Hermanowicz, Justyna Łabuz, Grażyna Dobrowolska, Olga Sztatelman

**Affiliations:** Institute of Biochemistry and Biophysics, Polish Academy of Sciences, Pawińskiego 5a, 02-106 Warszawa, Poland; Doctoral School of Molecular Biology and Biological Chemistry, Pawińskiego 5a, 02-106 Warszawa, Poland; Malopolska Centre of Biotechnology, Jagiellonian University, Gronostajowa 7A, 30-387 Kraków, Poland

## Abstract

Plants respond to changing environmental conditions through rapid cellular mechanisms, one of which is light-induced chloroplast movement. Plastid Movement Impaired 1 (PMI1) is one of the proteins involved in this process that undergoes rapid, blue light-dependent relocalization within the plasma membrane. Here, we investigate the molecular determinants of PMI1 membrane association. We identify the NT-C2 domain as the principal membrane-binding module and show that it extends beyond the boundaries previously assigned to the C2 domain. Plasma membrane localization is mediated by interactions between the extended NT-C2 domain and the phosphoinositides phosphatidylinositol 4-phosphate (PI4P) and phosphatidylinositol 4,5-bisphosphate [PI(4,5)P_2_], with basic residues within this region being essential for PMI1 membrane binding. We further demonstrate that the NT-C2 domain binds Ca^2+^ in vitro and that calcium availability modulates its phosphoinositide-binding preference. Consistently, depletion of cytosolic Ca^2+^ or inhibition of Ca^2+^ fluxes abolished the blue light-induced redistribution of PMI1 within the plasma membrane. Finally, we show that the extended NT-C2 domain, together with its flanking intrinsically disordered regions, is indispensable for PMI1 function in chloroplast movement regulation.

## Introduction

Chloroplast positioning in plant cells is dynamically regulated by light cues and serves as a mechanism for optimizing photosynthetic efficiency while minimizing photodamage. Under low-intensity light, chloroplasts redistribute to periclinal cell walls, perpendicular to the incident light, thereby maximizing light absorption (Gotoh et al., 2018; Zurzycki, 1955). In contrast, exposure to high-intensity light induces an avoidance response in which chloroplasts align along anticlinal walls parallel to the incoming light, reducing excessive light absorption and facilitating light penetration into deeper tissues (Davis et al., 2011; Kasahara et al., 2002; Sztatelman et al., 2010; Zurzycki, 1955). In higher plants, such as *Arabidopsis thaliana*, these chloroplast movement responses are triggered specifically by blue/UV wavelengths and are mediated by the phototropin family of photoreceptors (Briggs et al., 2001; Jarillo et al., 2001; Kagawa et al., 2001; Sakai et al., 2001). Arabidopsis contains two phototropins, phot1 and phot2, which act redundantly across multiple light-dependent processes—including phototropism, stomatal opening, leaf positioning, and chloroplast accumulation—with phot1 exhibiting greater sensitivity to low light (Briggs et al., 2001; Christie, 2007; de Carbonnel et al., 2010; Sakai et al., 2001).

Notably, only phot2 is capable of fully inducing the chloroplast avoidance response and additionally contributes to palisade cell development and chloroplast positioning in darkness (Jarillo et al., 2001; Kagawa et al., 2001; Suetsugu et al., 2005).

Phototropins localize primarily to the plasma membrane, with a fraction also associated with the chloroplast outer envelope (Kong et al., 2006; Kong, Suetsugu, et al., 2013; Sakamoto & Briggs, 2002). Plasma membrane localization is essential for mediating chloroplast accumulation, whereas chloroplast avoidance can be supported by chloroplast-envelope-localized phot2, emphasizing the importance of the chloroplast–plasma membrane interface in this process (Ishishita et al., 2020). Blue-light perception occurs through N-terminal LOV (light, oxygen, or voltage) domains, which in turn activate the C-terminal kinase domain, leading to photoreceptor autophosphorylation followed by the phosphorylation of downstream targets (for review see: Christie, 2007; Łabuz et al., 2022). Several proteins phosphorylated by phototropins that participate in different responses have been identified so far, but their targets involved in chloroplast relocations mostly await identification (Sullivan et al., 2021; Takemiya et al., 2013). Activation of phot1 and phot2 triggers a transient, minute-long increase in cytosolic Ca²⁺ following blue-light exposure. This rise results from Ca²⁺ influx through plasma membrane channels stimulated by both phototropins, while phot2 additionally promotes Ca²⁺ release from intracellular stores via phospholipase C-dependent inositol 1,4,5-triphosphate signaling (Harada et al. 2003). Evidence from inhibitor studies indicates that Ca²⁺ and phosphoinositide signaling contribute to the regulation of chloroplast movement (Aggarwal, Łabuz, et al., 2013, 2013; Anielska-Mazur et al., 2009; Grabalska & Malec, 2004; Tlałka & Fricker, 1999).

Although several downstream regulators have been identified, the signaling cascade linking blue-light perception to chloroplast movements has not been fully elucidated. Current evidence highlights a central role for the actin cytoskeleton, particularly short chloroplast-associated actin filaments (cp-actin) located between the plasma membrane and chloroplasts (Kadota et al., 2009). Upon blue-light exposure, cp-actin undergoes rapid reorganization: it disappears from the trailing edge while new filaments assemble at the leading edge. The polarized actin remodeling is proposed to provide the mechanical force that drives directional chloroplast movement (Kong, Arai, et al., 2013).

Several actin-associated proteins contribute to chloroplast positioning, and their function is closely linked to their localization (Banaś et al., 2012). CHLOROPLAST UNUSUAL POSITIONING 1 (CHUP1), which localizes to the chloroplast outer envelope, is required for anchoring chloroplasts to the plasma membrane, as *chup1* mutants exhibit chloroplast detachment and aggregation (Oikawa et al., 2003, 2008). CHUP1 interacts with actin and was recently identified as an actin polymerization factor (Kong et al., 2024). The kinesin-like proteins KINESIN-LIKE PROTEIN FOR ACTIN-BASED CHLOROPLAST MOVEMENT 1 and 2 (KAC1 and KAC2) also bind actin *in vitro* and are essential for the formation of cp-actin filaments surrounding chloroplasts; loss of KAC function results in impaired membrane attachment and defective chloroplast accumulation responses (Suetsugu et al., 2010). KAC1 additionally associates with the 14-3-3 proteins OMEGA and KAPPA (Dwyer & Hangarter, 2022).

THRUMIN1 is an actin-bundling protein that is tethered to the plasma membrane through N-terminal N-myristoylation and S-palmitoylation, a localization that is essential for its function (Whippo et al., 2011). THRUMIN1 associates with actin filaments in a light- and phototropin-dependent manner and localizes to cp-actin, dynamically redistributing to the leading edge of moving chloroplasts following irradiation. Although its actin-binding activity is regulated by phosphorylation, analysis of actin-binding–deficient THRUMIN1 variants revealed that chloroplast movement is largely preserved, suggesting that THRUMIN1 primarily functions as an anchor linking chloroplasts to the plasma membrane rather than as a direct driver of motility. Phosphorylation-dependent interaction of THRUMIN1 with 14-3-3 KAPPA and OMEGA proteins may further contribute to the assembly of chloroplast movement–associated protein complexes (Dwyer & Hangarter, 2021).

Members of the NON-PHOTOTROPIC HYPOCOTYL 3 (NPH3)/ROOT PHOTOTROPISM 2 (RPT2)-LIKE (NRL) protein family also interact with 14-3-3 proteins and act as phototropin substrates with distinct functional specificities (Sullivan et al., 2021). RPT2 and NRL PROTEIN FOR CHLOROPLAST MOVEMENT 1 (NCH1) redundantly mediate the chloroplast accumulation response but are not required for avoidance (Suetsugu et al., 2016). RPT2 is phosphorylated by phototropins at a conserved C-terminal serine residue (S591), which promotes its association with 14-3-3 proteins and is essential for its role in chloroplast accumulation (Waksman et al., 2023).

Additional regulators of cp-actin dynamics include J-DOMAIN PROTEIN REQUIRED FOR CHLOROPLAST ACCUMULATION RESPONSE 1 (JAC1), WEAK CHLOROPLAST MOVEMENT UNDER BLUE LIGHT 1 (WEB1), and PLASTID MOVEMENT IMPAIRED 2 (PMI2), which function as a complex (Kodama et al., 2010; Luesse et al., 2006; Suetsugu et al., 2005). JAC1 is specifically required for the accumulation response, whereas *web1* and *pmi2* mutants display attenuated, but not abolished, chloroplast movement. WEB1 and PMI2 are proposed to suppress JAC1-mediated accumulation under high-light conditions, and mutations in these proteins alter cp-actin sensitivity to blue light (Kodama et al., 2010). Collectively, these factors are thought to modulate cp-actin dynamics rather than directly drive actin polymerization (Wada and Kong, 2018).

Plastid Movement Impaired 1 (PMI1) is another key regulator of chloroplast movement. PMI1 is an 843-amino acid protein with two predicted conserved domains—an N-terminal C2 domain (NT-C2, residues 133–288) and a C-terminal domain of unknown function (C-DUF, beginning at residue 517), separated by two intrinsically disordered regions (DeBlasio et al., 2005). PMI1 is a plasma membrane-bound protein that dynamically redistributes in response to changes in light conditions: after dark acclimation, it localizes to cortical and cp-actin filaments, but under blue-light illumination of only part of the cell, it relocates from lit to shaded membrane regions. This movement depends on phototropins and occurs even when

F-actin is disrupted, indicating that PMI1 responds upstream of actin remodeling. PMI1 forms a light-dependent complex with THRUMIN1 and KAC1 via the 14-3-3 KAPPA and OMEGA proteins, with only the PMI1–14-3-3 interaction being directly regulated by light. These observations support a model in which PMI1 displacement promotes chloroplast de-anchoring, enabling it to move, whereas its re-association in shaded regions facilitates re-anchoring during repositioning (Dwyer & Hangarter, 2022).

PMI1 function depends on its plasma membrane localization, yet the mechanism underlying this association remains unclear. The presence of a C2 domain suggests a lipid-binding role, as C2 domains are common in eukaryotic proteins and mediate membrane association, often in a Ca^2+^-dependent manner (Cho & Stahelin, 2006). Phylogenetic analysis places the PMI1 C2 domain in the NT-C2 subfamily, characterized by an N-terminal C2 domain followed by regions involved in cytoskeletal interactions (Zhang & Aravind, 2010).

Here, we identify the region responsible for PMI1 plasma membrane localization, determine the lipids it recognizes, and examine the role of calcium in these interactions. Finally, using mutants with defined deletions, we show that the lipid-binding module, together with its flanking regions are essential for chloroplast relocation.

## Results

### PMI1 Membrane Association Requires an Extended NT-C2 Region

To identify the regions responsible for membrane association, we first examined the intracellular distribution of fluorescently tagged PMI1 and its fragments transiently expressed in *Arabidopsis thaliana* leaves. PMI1 was divided into four parts—an N-terminal unstructured region (aa 1–132), the predicted core NT-C2 domain (aa 133–288), a middle unstructured region (aa 289–516), and a C-terminal DUF (aa 517–843), based on (Zhang & Aravind, 2010) and (DeBlasio et al., 2005). None of the individual fragments, in contrast to the full-length PMI1, localized to the plasma membrane. Instead, all appeared in the cytoplasm; the N-terminal region and NT-C2 domain could be also detected in the nucleus, and the C-terminal DUF was observed in additional globular structures (Figure 1A). Thus, membrane affinity was lost upon fragmentation, and the predicted core NT-C2 domain alone was insufficient for membrane binding.

**Figure 1.**
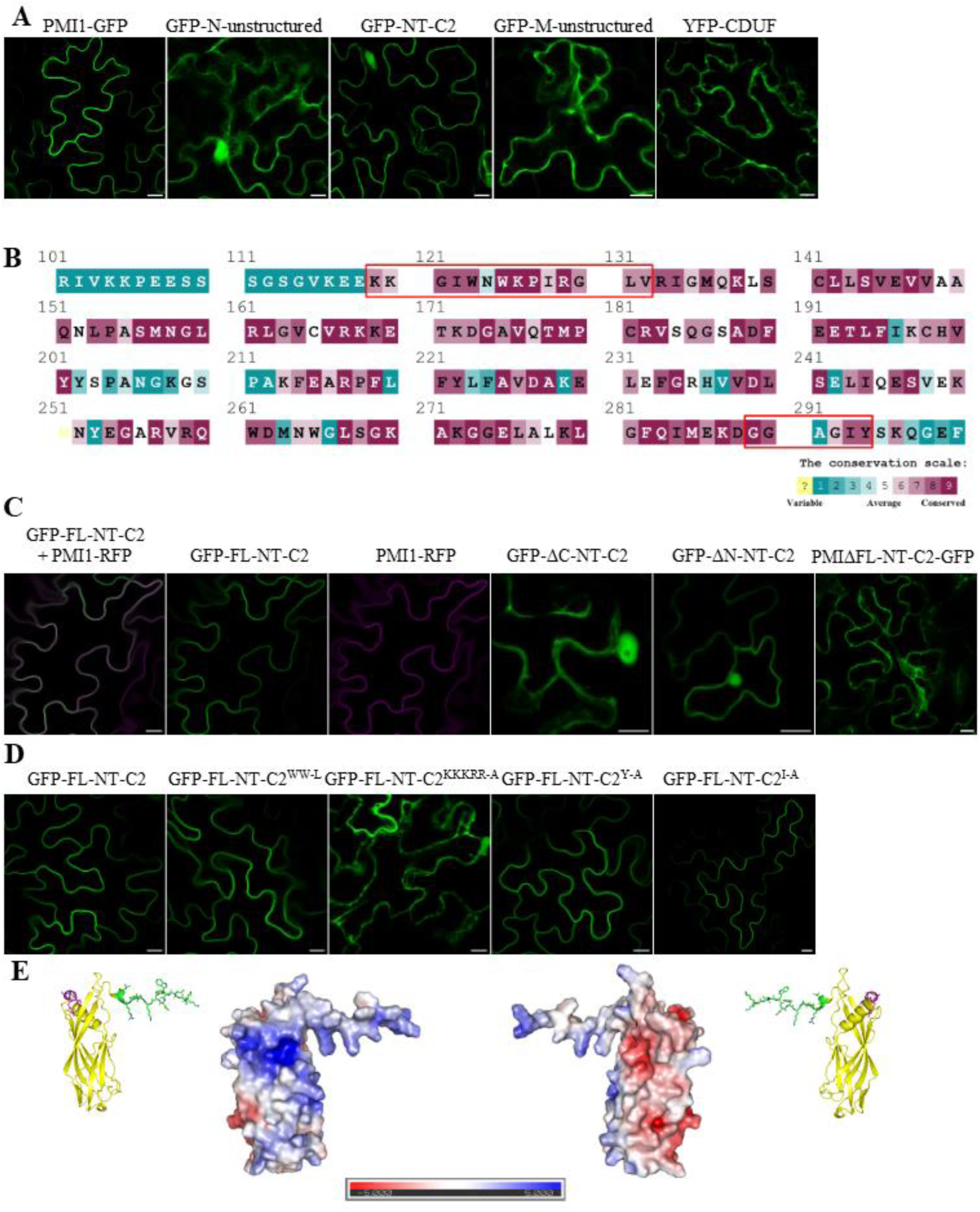
**Mapping of the PMI1 domain responsible for plasma membrane localization.** (A) Confocal images of GFP-tagged PMI1 and its fragments transiently expressed in *Arabidopsis thaliana* leaf epidermal cells. Full-length PMI1 localizes to the plasma membrane, whereas the N-terminal unstructured region, the core NT-C2 domain, the M-unstructured region, and the C-terminal DUF domain show predominantly cytoplasmic localization; the two N-terminal fragments additionally accumulate in the nucleus (scale bar = 10 μm). Microscopic images are representative of at least three independent biological replicates. (B) Conservation analysis of the PMI1 region encompassing the core NT-C2 domain (residues 133–288) based on ConSurf predictions (Ashkenazy et al., 2016). Red boxes indicate conserved N- and C-terminal extensions excluded from the core construct but included in the FL-NT-C2 variant. (C) FL-NT-C2 localizes to the plasma membrane in *A. thaliana* epidermal cells and co-localizes with full-length PMI1 when co-expressed. Deletion of either the C-terminal (GFP-ΔC-NT-C2) or N-terminal (GFP-ΔN-NT-C2) conserved extension abolishes plasma membrane localization, similar to PMI1 lacking the FL-NT-C2 region (PMI1ΔN-NT-C2-GFP) (scale bar = 10 μm). (D) Confocal images of GFP-tagged FL-NT-C2 point mutants transiently expressed in *A. thaliana* leaf epidermis. The WW-L (W123L, W125L), Y-A (Y294A), and I-A (I293A) variants retain plasma membrane localization comparable to the wild-type protein, whereas the KKKRR-A mutant (K119A, K120A, K126A, R129A, R133A) is cytoplasmic (scale bar = 10 μm). (E) Electrostatic surface representation of the PMI1 FL-NT-C2 domain generated in PyMOL (v2.5.2) using an AlphaFold2-predicted structure (Jumper et al., 2021). The positively charged surface likely faces the plasma membrane and mediates electrostatic interactions with anionic lipids, whereas the opposite, negatively charged surface is oriented toward the cytoplasm. Core NT-C2 is shown in yellow, with the N-terminal extension in green and the C-terminal extension in magenta.

In order to better understand the domain architecture of PMI1, we performed a homology search using the Consurf server (Ashkenazy et al., 2016). In contrast to previous reports based on homology with other C2 domains present among all eucaryotes (Zhang & Aravind, 2010), only plant-specific proteins with PMI1 homology were assessed. We found that the conserved region in PMI1 homologs is longer than expected and that at both the N-terminus and C-terminus of the core NT-C2 domain, there are stretches of highly conserved amino acids (Figure 1B, red boxes), and the whole conserved region spans from amino acids 119 to 294. When expressed *in planta*, this extended region, designated FL-NT-C2, localized to the plasma membrane and fully matched the localization of the full-length protein when co-expressed (Figure 1C). Both flanking sequences proved necessary for membrane targeting: N- or C-terminal truncation abolished membrane localization, producing a cytoplasmic signal indistinguishable from the core NT-C2 alone (Figure 1C).

Consistent with this requirement, deleting the FL-NT-C2 region from full-length PMI1 eliminated membrane binding entirely (Figure 1C).

### Electrostatic Interactions Drive FL-NT-C2 Membrane Association

To identify residues contributing to membrane interactions, we generated and expressed a series of point-mutant versions of FL-NT-C2 in *Arabidopsis* leaves. Mutations targeted aromatic residues (W123, W125, Y294), hydrophobic residue (I293), and clusters of basic residues (K119, K120, K126, R129, R133). Surprisingly, all mutants except the basic-residue mutant retained normal plasma-membrane localization. Only the KKKRR to AAAAA substitution caused complete loss of membrane association (Figure 1D). These results indicate that membrane binding does not rely on individual aromatic or hydrophobic residues but instead depends on a positively charged surface patch.

The structure of the PMI1 protein was not experimentally established, but structural modeling using AlphaFold2, followed by electrostatic surface representation, showed a pronounced polarization of the FL-NT-C2 domain, with a strongly positive surface opposite a negatively charged region (Figure 1E). The requirement for multiple basic residues is consistent with electrostatic interactions between FL-NT-C2 and anionic lipids in the plasma membrane.

### Extended NT-C2 Region Binds Phosphoinositides In Vitro

To assess lipid binding properties of the NT-C2 domain *in vitro*, FL-NT-C2 as well as truncated versions were purified as GST-fusions from *E. coli.* The Protein-Lipid Overlay (PLO) assay was performed according to Shirey et al. (2017) using membranes containing a broad range of biologically relevant phospholipids: Membrane Lipid Strips (Echelon Biosciences). The assay showed that the FL-NT-C2 domain interacts with PIPs: PI4P, PI(4,5)P2, and PI(3,4,5)P3 with varying affinities.

The signal was the strongest for monophosphate, and it was decreasing with every other phosphate group present at the inositol ring of the molecule. In the case of core NT-C2, ΔC-NT-C2, and ΔN-NT-C2, the residual signal was noticeable only for PI4P or was not visible at all (Figure 2A).

**Figure 2.**
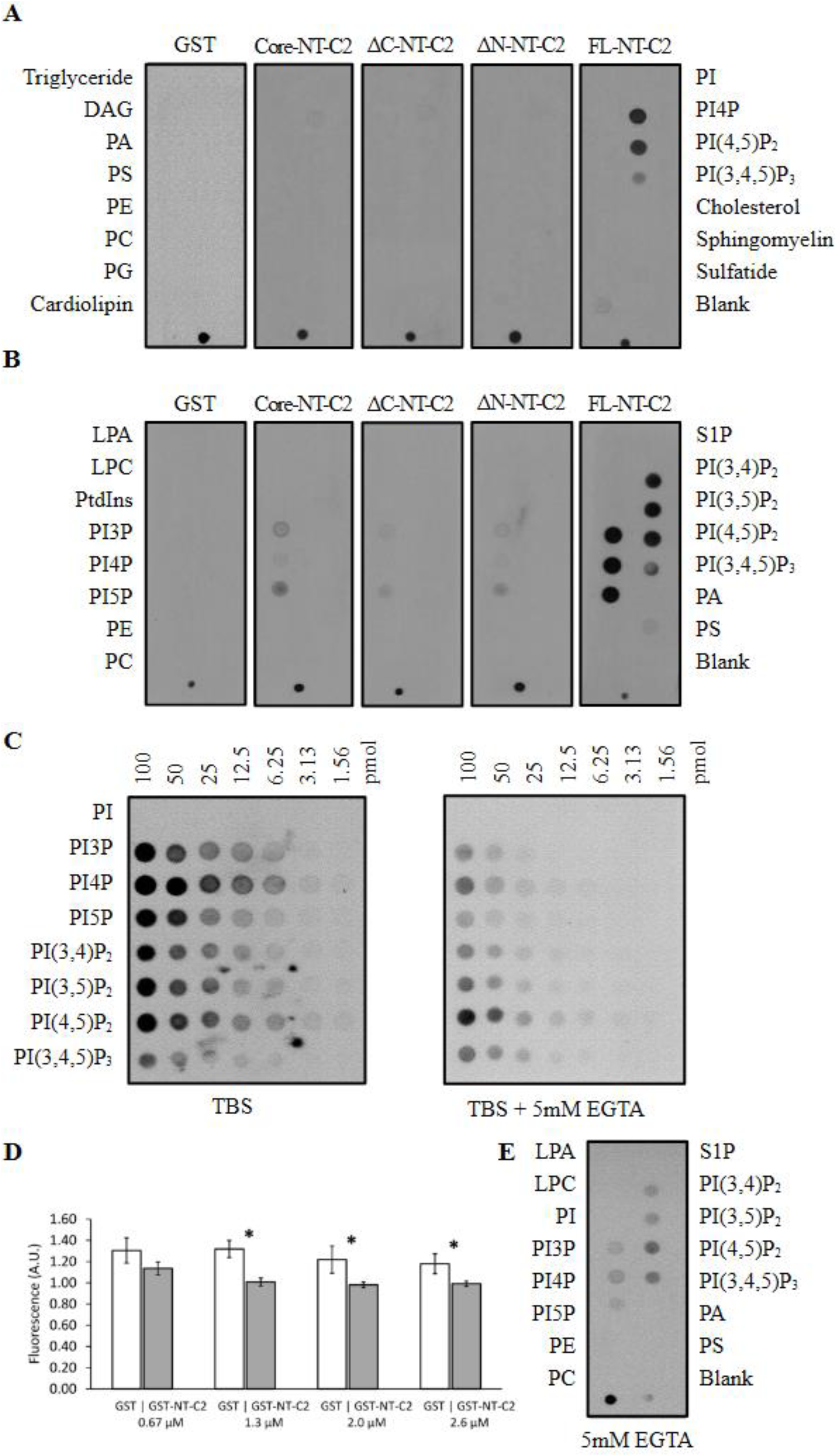
**In vitro lipid preference of the PMI1-NT-C2 domain and the influence of calcium.**-----(A, B) GST-tagged recombinant proteins were analyzed by protein–lipid overlay (PLO) assays using Membrane Lipid Strips (A) and PIP Strips (B) (Echelon Biosciences), respectively. The FL-NT-C2 domain binds phosphoinositides, with binding strength decreasing as the number of phosphate groups on the inositol headgroup increases. In contrast, the core NT-C2, ΔC-NT-C2, and ΔN-NT-C2 variants display only weak binding to phosphatidylinositol monophosphates.-----(C) PIP Array membranes (Echelon Biosciences) containing graded amounts of lipids were incubated with GST-tagged FL-NT-C2. Strong binding to PI4P and PI(4,5)P₂ was detected even at the lowest lipid concentration (1.56 pmol per spot). The addition of EGTA to the incubation buffer reduced binding to phosphatidylinositol monophosphates.-----(D) Calcium binding by the FL-NT-C2 domain was assessed using the fluorescent dye Fura-2. GST-tagged FL-NT-C2 and GST alone (control) were pretreated with EGTA, desalted, and incubated at equimolar concentrations (0.67–2.6 μM) with 33 μM Ca²⁺ and glutathione Sepharose. Free calcium in the supernatant was quantified by Fura-2 fluorescence (excitation 350 nm, emission 506 nm). Significantly lower fluorescence in FL-NT-C2 samples compared with GST controls indicates calcium binding by the PMI1 C2 domain (Student’s *t*-test with Bonferroni correction for four comparisons (α = 0.0125; *p* = 0.022, <0.001, 0.004, 0.003).-----(E) PLO assays using PIP Strips in the presence of 5 mM EGTA showed reduced binding of FL-NT-C2 to phosphatidylinositol monophosphates in the absence of Ca²⁺, whereas binding to PI(4,5)P₂ remained relatively strong.-----For all PLO assays (A–C, E), membranes were blocked with TBS-T containing 3% BSA (with 5 mM EGTA where indicated) and incubated with 0.5 μg/mL protein. Anti-GST primary antibodies were used for detection, and all protein and antibody solutions were prepared in blocking buffers. Data shown are representative of three independent replicates.

Subsequently, lipid-binding assays were performed using membranes containing a broad range of phosphoinositides (PIP Strips; Echelon Biosciences). The FL-NT-C2 domain bound all PIPs, with strongest binding to PI3P, PI4P, and PI5P; weaker binding to PI(3,4)P2, PI(3,5)P2, and PI(4,5)P2; and the weakest binding to PI(3,4,5)P3. In contrast, the core NT-C2 domain and the truncated ΔN-NT-C2 and ΔC-NT-C2 variants showed no binding to phosphatidylinositol bis- or trisphosphates and reduced binding to monophosphates compared with FL-NT-C2 (Figure 2B).

To further resolve differences in binding affinity, PIP Array membranes containing graded amounts of PIPs (100 to 1.56 pmol per spot) were used. The strongest binding, detectable even at the lowest lipid concentrations, was observed for PI4P and PI(4,5)P2, whereas binding to other PIPs was weaker (Figure 2C).

### FL-NT-C2 Binds Calcium and Exhibits Ca²⁺-Dependent Lipid Selectivity

Calcium ions can modulate the function of C2 domains, either enabling lipid binding or altering membrane-binding affinity (Cho & Stahelin, 2006). Therefore, we examined whether FL-NT-C2 binds Ca^2+^ in vitro. A Fura-2 fluorescence assay revealed significantly greater Ca^2+^ binding by FL-NT-C2 compared with GST alone at higher concentrations of the protein (Figure 2D).

Next, we tested whether Ca^2+^ affects lipid binding. In standard assays, performed without added Ca^2+^ but lacking chelators, FL-NT-C2 strongly preferred monophosphorylated PIPs. However, EGTA-mediated Ca^2+^ depletion shifted lipid specificity: binding to monophosphates diminished, while binding to more highly phosphorylated species increased, with PI(4,5)P2 becoming the dominant target (Figure 2C, E). These findings indicate that Ca^2+^ modulates FL-NT-C2 lipid selectivity, with PI(4,5)P2 binding being less Ca^2+^-dependent than PI4P binding.

### FL-NT-C2 Associates with PI4P and PI(4,5)P2 In Planta

To validate lipid interactions *in vivo*, we transiently expressed RFP-tagged FL-NT-C2 in *Arabidopsis* PIP-line plants expressing well-characterized phosphoinositide markers (Simon et al., 2014). FL-NT-C2 co-localized with the PI4P marker 2xPH(FAPP1) and the PI(4,5)P2 marker 2xPH(PLC), but did not co-localize with the PI3P marker (Figure 3A–C).

**Figure 3.**
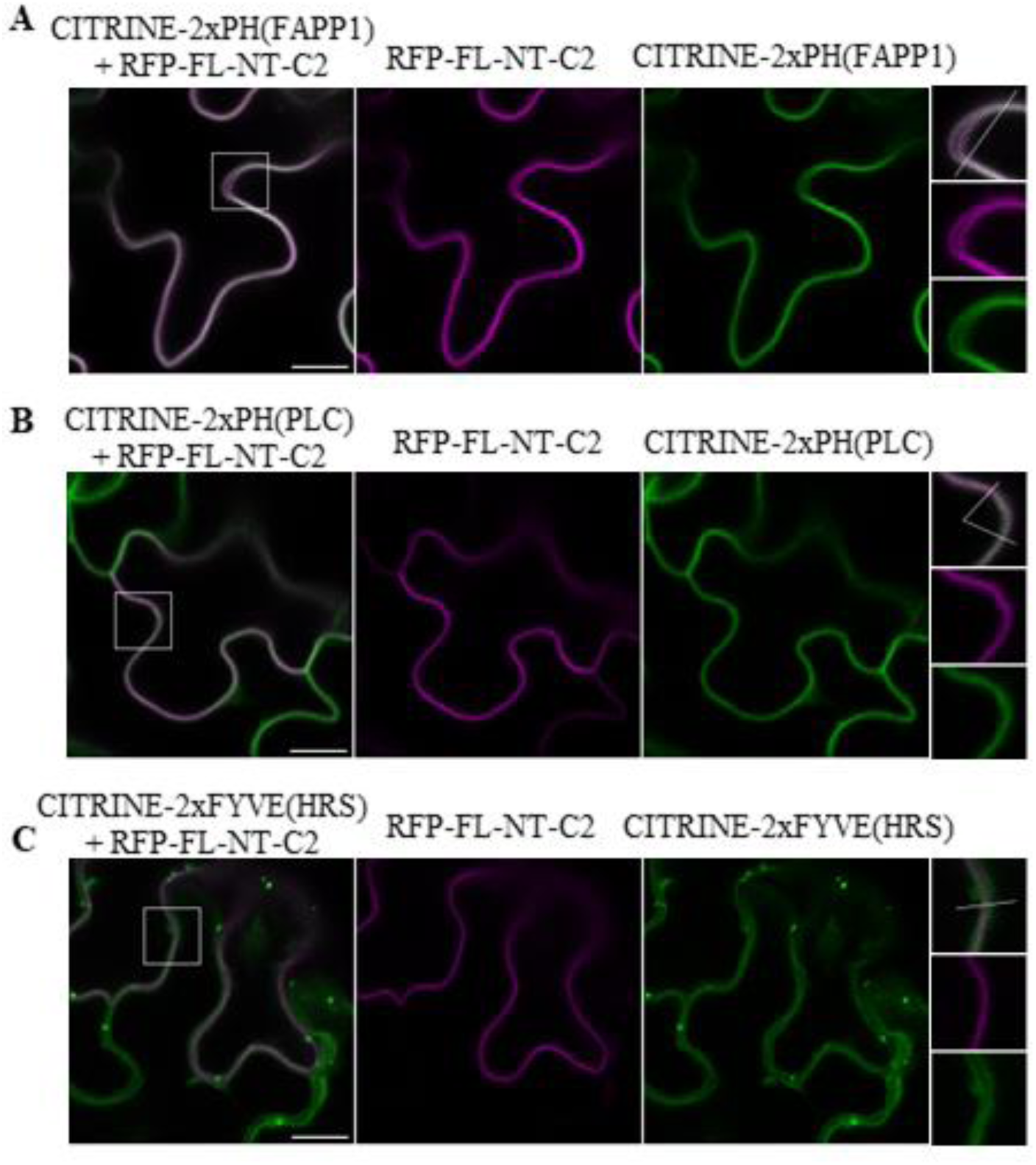
**Co-localization of PMI1-FL-NT-C2 with phosphoinositide markers in Arabidopsis.** RFP-tagged FL-NT-C2 was transiently expressed in leaves of *Arabidopsis thaliana* PIP-line plants (Simon et al., 2014) expressing CITRINE-tagged phosphoinositide markers for (A) PI4P, (B) PI(4,5)P2, and (C) PI3P. FL-NT-C2 co-localizes with the PI4P marker 2xPH(FAPP1) and the PI(4,5)P2 marker 2xPH(PLC), but shows no co-localization with the PI3P marker 2xFYVE(HRS). Boxed regions are shown as magnified views on the right. Scale bar = 10 μm. Data shown are representative of at least three independent biological replicates.

Several *in planta* approaches were employed to discriminate between PI4P and PI(4,5)P2 binding by FL-NT-C2. Phenylarsine oxide (PAO), a PI4K inhibitor ^(^Wiedemann et al., 1996^)^, was first tested based on reports that short-term treatment depletes plasma membrane PI4P without affecting PI(4,5)P₂ in roots (Simon et al., 2016). However, in leaves of PIP-line plants expressing the PI4P marker 2xPH(FAPP1), treatment with 60 μM PAO did not alter marker localization, even after prolonged incubation or increased inhibitor concentration (Supplementary Figure S1), indicating that PAO is ineffective in leaf tissue, likely due to detoxification mechanisms.

To overcome this limitation, PAO was applied to roots, where stable 35S::PMI1-YFP lines allow ectopic PMI1 expression. In response to PAO treatment, as expected, PI4P marker localization shifted from the plasma membrane to the cytoplasm, whereas the PI(4,5)P2 marker was largely unaffected (Figure 4A). Occasional relocalization of the PI(4,5)P2 marker was also observed. PMI1-YFP relocalized from the plasma membrane, although with somewhat variable timing and spatial heterogeneity. These results suggest that PMI1 interacts with PI4P, PI(4,5)P2, or both; however, because PI4P depletion also impacts PI(4,5)P2 synthesis, this approach did not clearly distinguish between the two lipid pools.

**Figure 4.**
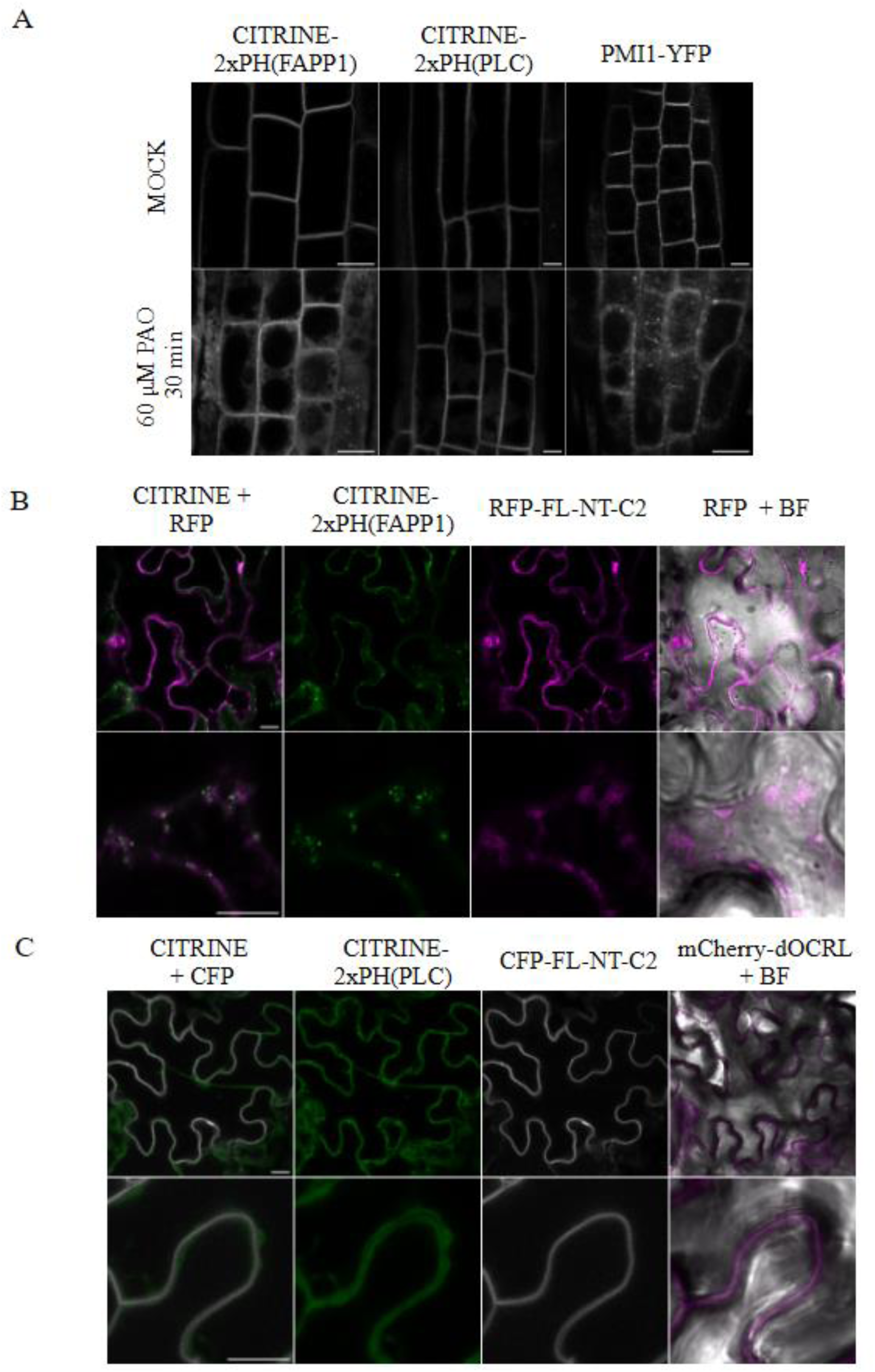
**Effects of PI4P and PI(4,5)P₂ depletion on PMI1 or PMI1-NT-C2 localization.** (A) In root cells, PMI1 relocalizes from the plasma membrane upon PAO treatment. After 30 min, a minor relocalization of the PI(4,5)P2 marker CITRINE–2xPH(PLC) is also observed. Compared with the PI4P marker CITRINE–2xPH(FAPP1), PMI1 shows a weaker and more heterogeneous response, varying among individual cells. (B) Treatment with 30 μM wortmannin for 90 min causes partial dissociation of the PI4P marker CITRINE–2xPH(FAPP1) from the plasma membrane and its accumulation in numerous cytoplasmic puncta. In contrast, RFP–FL-NT-C2 becomes more diffuse at the cell periphery, with occasional cytoplasmic strands, but does not accumulate in the same punctate structures. (C) In the iDePP system, 48 h after dexamethasone (DEX) induction, the PI(4,5)P₂ marker CITRINE–2xPH(PLC) relocalizes to the cytoplasm, whereas the FL-NT-C2 domain remains associated with the plasma membrane and does not co-relocate with the marker. Bright-field images in panels (B) and (C) were adjusted for clarity using the Brightness/Contrast function in ImageJ. Data shown are representative of at least six independent biological replicates. Scale bar = 10 μm

Wortmannin (WM), a known inhibitor of PI3K (Delage et al., 2012; Jaillais et al., 2006), which at higher concentrations (>30 μM) inhibits PI4K, was therefore used as an alternative, as its effect in leaves was reported previously (Anielska-Mazur et al., 2009). RFP-FL-NT-C2 was transiently expressed in leaves of PIP-line plants, which were subsequently treated with 30 μM WM for 90 min. The PI4P marker CITRINE-2xPH(FAPP1) partially dissociated from the plasma membrane and accumulated in cytoplasmic puncta in response to the treatment (Figure 4B). In contrast, FL-NT-C2 did not relocalize to these structures but instead showed increased diffusion at the plasma membrane and occasional cytoplasmic strands, indicating a distinct response to PI4K inhibition.

Finally, PI(4,5)P2 specificity was directly tested using the iDePP system (Doumane et al., 2021), which enables inducible depletion of PI(4,5)P2 without affecting PI4P. CFP-FL-NT-C2 was transiently expressed in plants carrying DEX-inducible mCherry-dOCRL (PI(4,5)P₂ phosphatase) and the PI(4,5)P2 marker CITRINE-2xPH(PLC).

Following DEX induction, the PI(4,5)P2 marker was displaced from the plasma membrane, whereas FL-NT-C2 remained membrane-associated (Figure 4C), indicating that PI(4,5)P2 is not the primary determinant of FL-NT-C2 plasma membrane localization.

Taken together, these experiments indicate that PMI1 can interact with both PI4P and PI(4,5)P2, but PI4P depletion affects its membrane association more strongly.

### Calcium Is Required for PMI1 Light-Induced Redistribution

To assess the role of calcium in PMI1 localization and light-induced redistribution, Arabidopsis plants stably expressing 35S::PMI1-YFP were analyzed. Dark-adapted detached leaves were infiltrated with A23187 plus EGTA to deplete intracellular calcium or with A23187 plus Ca²⁺ to promote calcium influx, incubated in darkness, and imaged by confocal microscopy at the cortical plane of palisade mesophyll cells using 514 nm excitation.

Calcium depletion caused a clear change in PMI1 distribution, from a uniform plasma membrane pattern to filamentous membrane-associated structures, whereas elevated calcium had no effect on dark-state localization (Figure 5A, B, darkness).

**Figure 5.**
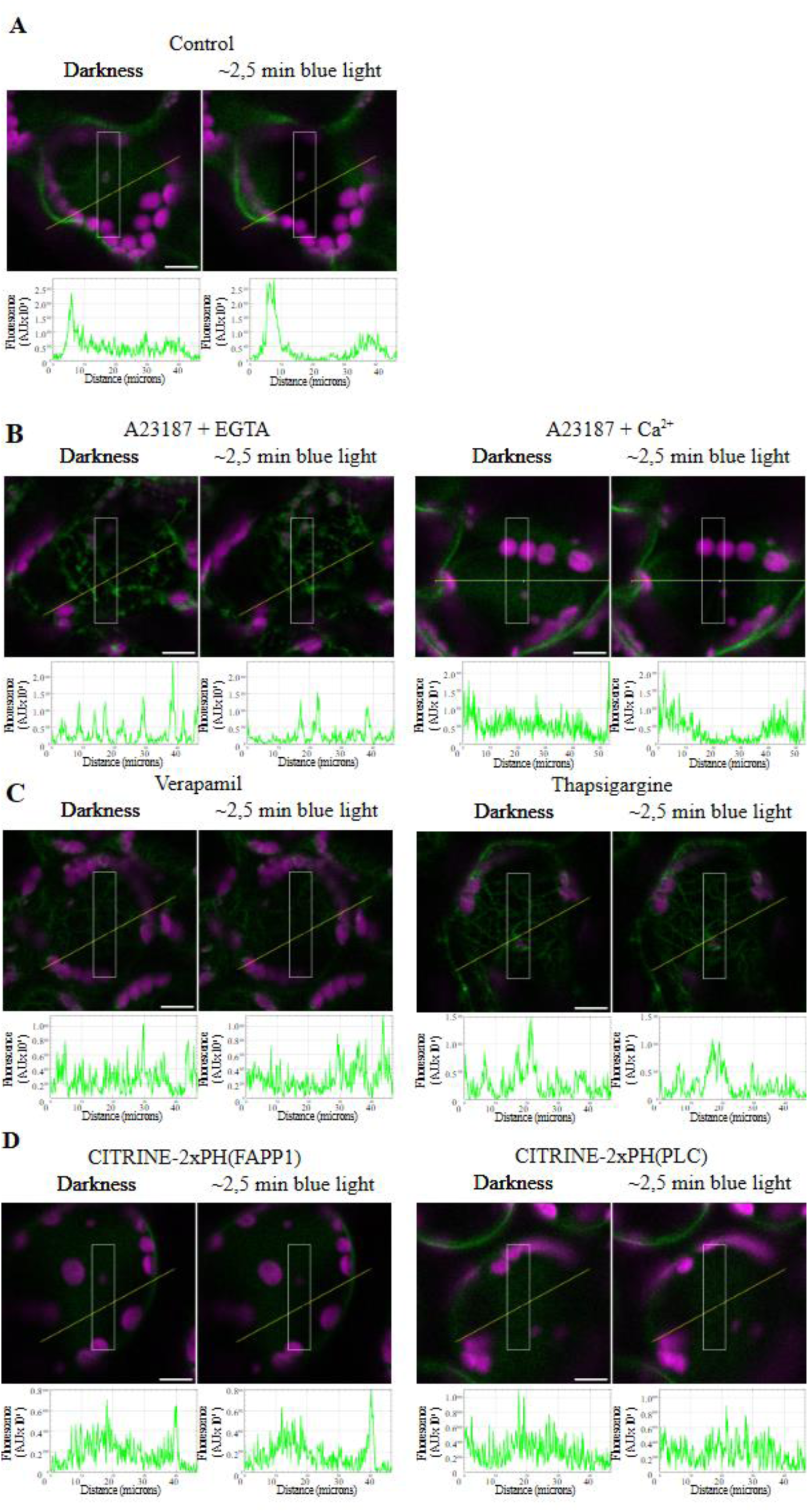
**Effect of calcium modulation on PMI1 localization in planta.** Arabidopsis plants expressing PMI1-YFP were treated with compounds altering calcium levels, and a selected plasma membrane region (white box) was irradiated with a 485 nm laser for 2.5 min. (A) Control conditions. (B) Treatment with 10 µM A23187 + 1 mM EGTA caused PMI1 to shift from a uniform plasma membrane distribution to filamentous membrane-associated structures, preventing redistribution to the unirradiated region under blue light. 10 µM A23187 + 10 mM Ca²⁺ had no effect on localization or light-induced movement. (C) 20 µM verapamil or 15 µM thapsigargin treatments mimicked the effect of EGTA, inducing filamentous PMI1 distribution and blocking redistribution. (D) PI4P (CITRINE-2xPH(FAPP1)) and PI(4,5)P2 (CITRINE-2xPH(PLC)) markers did not relocalize upon blue-light exposure. Fluorescence intensity profiles were generated along the yellow lines in the images using the Plot Profile function in ImageJ. Data shown are representative of at least six independent biological replicates.

To test light-induced redistribution, a defined plasma membrane region was illuminated with 458 nm light for ∼2.5 min based on Dwyer & Hangarter (2022). Under control and A23187 + Ca²⁺ conditions, PMI1 relocated away from the irradiated area, but this response was completely abolished after EGTA treatment, with PMI1 remaining in filamentous structures despite blue-light exposure (Figure 5B).

To distinguish between extracellular calcium influx and intracellular calcium stores, plants were treated with the calcium channel blocker verapamil or the intracellular store-depleting agent thapsigargin. Both treatments phenocopied calcium depletion, resulting in filamentous PMI1 localization and loss of light-induced displacement (Figure 5C), indicating that membrane-associated PMI1 and calcium availability are required for the redistribution process.

Finally, to test whether PMI1 movement reflects phosphoinositide redistribution within the plasma membrane, PI4P and PI(4,5)P2 markers were examined following blue-light irradiation. Neither CITRINE–2xPH(FAPP1) nor CITRINE–2xPH(PLC) showed detectable relocalization (Figure 5D), indicating that PMI1 displacement is not driven by bulk movement of these lipids within the plasma membrane.

### PMI1 Unstructured Regions and the NT-C2 Domain are Required for Chloroplast Movements

To examine the relationship between the domain architecture of PMI1 and protein function, we used CRISPR–Cas9 to generate a series of *Arabidopsis thaliana* lines carrying truncations in the *PMI1* gene. Guide RNAs were designed to target the boundaries of predicted PMI1 regions and paired with a guide targeting the gene terminus, yielding a set of C-terminal truncation lines ranging from deletion of the C-DUF domain alone, through progressive loss of the middle unstructured region and the NT-C2 domain, to complete *pmi1* knockouts.

In total, 14 independent lines were characterized (Figure 6A). The first digit in each line identifier denotes the gRNA pair used for its generation. For most lines, sequencing confirmed deletions consistent with the predicted Cas9 cut sites and removal of the intervening genomic region, although several lines exhibited larger deletions or more complex genomic rearrangements. Line 1.8.4 carries a deletion encompassing the entire M-unstructured region and part of the C-DUF, whereas line 1.1.15, generated with the same gRNA pair, retains the N-unstructured region and NT-C2 domain but lacks the C-terminal part of PMI1. Lines 5.12.12.1 and 5.7.12 lack only the C-DUF region, while line 5.12.12.3 carries a larger deletion and is predicted to encode only the N-unstructured region and NT-C2 domain. In line 6.18.30, an additional inverted DNA fragment was inserted at the 5′ Cas9 cut site, unlike line 6.1.2. Lines 7.6.4 and 7.8.8.6 retain only a fragment of the N-unstructured region. Lines 4.20.15.1 and those with identifiers starting with 8 or 9 represent complete *pmi1* knockouts.

**Figure 6.**
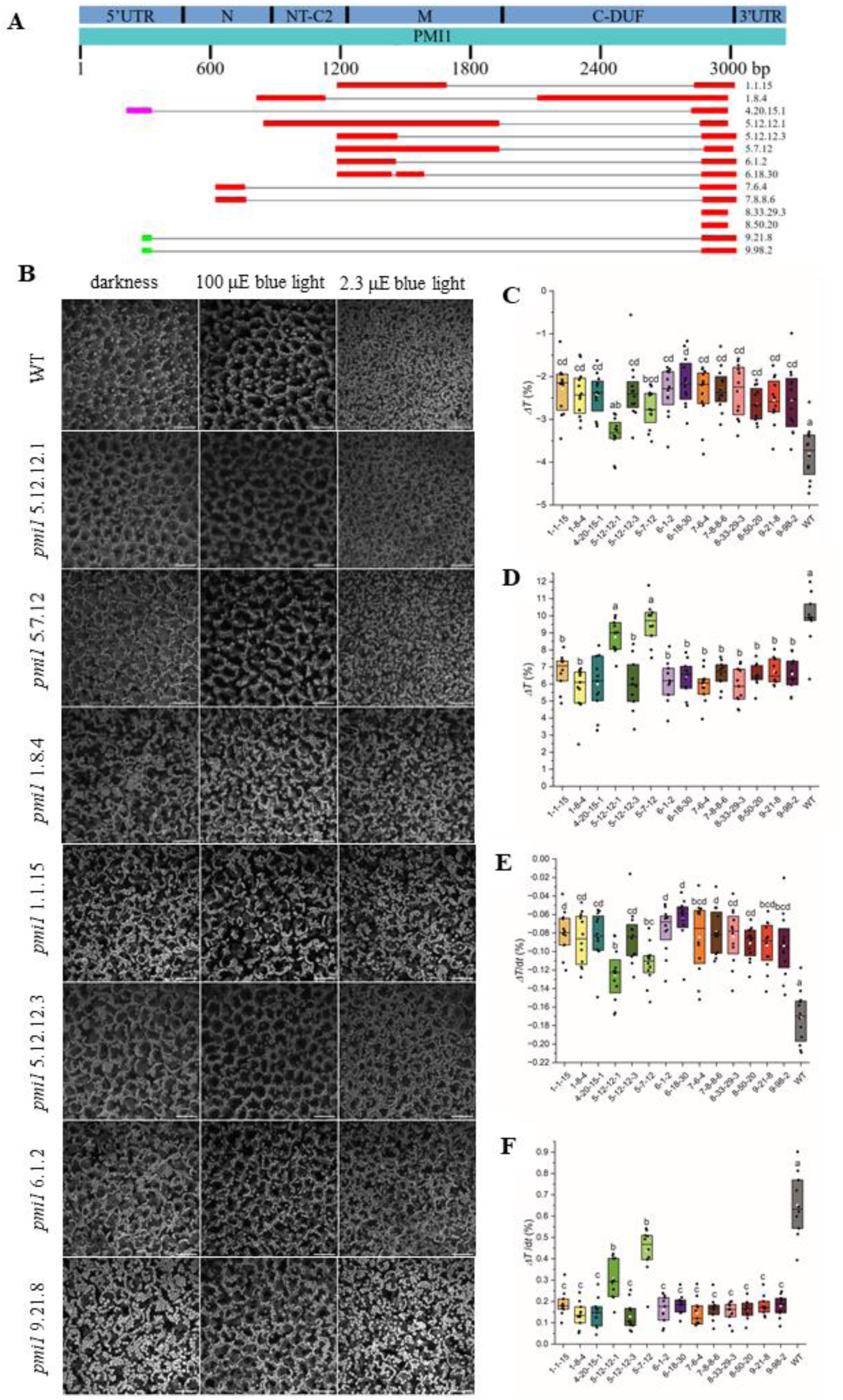
**Characterization of *pmi1* CRISPR–Cas9 mutant lines and their chloroplast movement phenotypes.** (A) Comparison of nucleotide sequences of *pmi1* mutant alleles generated by CRISPR–Cas9, visualized using NCBI BLAST and aligned to the wild-type *Arabidopsis thaliana* PMI1 sequence. Boxes indicate aligned regions, while grey lines or their absence (lines 8.33.29.3 and 8.50.20) denote deletions. N and M indicate the N-terminal and middle unstructured regions, respectively. (B) Chloroplast movements in wild-type and selected *pmi1* mutant plants visualized by confocal microscopy using chlorophyll autofluorescence. In wild type, chloroplasts are evenly distributed along the cell bottom and partially along anticlinal walls in darkness, relocate to anticlinal walls under high-intensity blue light (100 μmol m^−2^ s^-1^, 1 h; avoidance response), and accumulate beneath the upper periclinal wall under low-intensity blue light (2.3 μmol m^−2^ s^-1^; accumulation response). Most *pmi1* mutants show abnormal chloroplast positioning already in darkness, with clustered chloroplasts resembling the accumulation state and a reduced avoidance response. In contrast, lines *5.12.12.1* and *5.7.12* display wild-type–like chloroplast positioning and intact accumulation and avoidance responses (scale bar = 50 μm). Images shown are representative of ten independent biological replicates. (C–F) Quantitative analysis of chloroplast movements in wild type and *pmi1* mutants based on red light transmittance measurements. (C, D) Response amplitudes and (E, F) response velocities to (C, E) low-intensity blue light (1 μmol m^−2^ s^-1^; accumulation) and (D, F) high-intensity blue light (100 μmol m^−2^ s^-1^; avoidance). Most *pmi1* mutants exhibit reduced amplitudes and velocities compared with wild type, whereas lines *5.12.12.1* and *5.7.12* show amplitudes comparable to wild type and intermediate reaction velocities. 10-12 leaves were measured for each group. The white square represents the mean, the black horizontal line represents the median, and the box represents the 25-75 percentile. Each dot shows one measured leaf. Means of groups that do not share a common letter are different at a 0.05 significance level as tested with Tukey’s test.

Chloroplast movements in mutant plants were analyzed by confocal microscopy using chlorophyll autofluorescence. Leaves from dark-adapted plants were exposed to blue light, and chloroplast positioning was assessed through z-stack imaging of palisade mesophyll cells. In wild-type plants, chloroplasts were evenly distributed along the cell bottom and partially along anticlinal walls in darkness, accumulated along anticlinal walls under high-intensity blue light (100 µmol m^−2^ s^-1^), and relocated beneath the upper periclinal wall under low-intensity blue light (2.3 µmol m^−2^ s^-1^) (Figure 6B; Supplementary Figure S2). In contrast, most mutant lines exhibited abnormal chloroplast positioning already in darkness, with chloroplasts pre-accumulated beneath the upper cell wall, and showed a strongly impaired avoidance response under high-intensity light. Notably, two mutant lines, *5.12.12.1* and *5.7.12*, displayed wild-type–like chloroplast distribution in darkness as well as intact accumulation and avoidance responses (Figure 6B). These lines uniquely retain an intact coding sequence for the middle unstructured region and upstream domains of PMI1, suggesting that these regions are required for PMI1 function in chloroplast movement.

To quantify these phenotypes, leaf transmittance assays were performed. Under low-intensity blue light (1 µmol m^−2^ s^-1^), wild-type leaves showed a decrease in transmittance consistent with chloroplast accumulation, whereas most *pmi1* mutants displayed reduced responses. The *pmi1 5.12.12.1* line behaved similarly to wild type, while *pmi1 5.7.12* showed an intermediate response (Figure 6C). Under high-intensity blue light (100 µmol m^−2^ s^-1^), transmittance increased in wild type, indicating chloroplast avoidance; again, most mutants showed attenuated responses, with *pmi1 5.12.12.1* and *pmi1 5.7.12* statistically indistinguishable from wild type (Figure 6D). Similar trends were observed in response kinetics (Figure 6E, F).

## Discussion

PMI1 is anchored to the plasma membrane through its NT-C2 domain, which binds phosphoinositides. Our data demonstrate that this domain is larger than previously predicted and includes additional N- and C-terminal extensions. Both flanking regions are required for efficient plasma membrane localization *in vivo* and for lipid binding *in vitro*. While a similar domain architecture appears conserved among plant proteins, it has not been described previously for eukaryotic NT-C2 domains, suggesting that plant-specific NT-C2 domains may employ distinct mechanisms of membrane lipid interaction. Notably, unlike several other NT-C2–containing proteins in which this domain is located at the extreme N terminus, PMI1 contains an unstructured N-terminal region preceding the NT-C2 domain. To explore the potential function of this region, we performed computational disorder predictions using IUPred3 (Erdős et al., 2021) and ANCHOR2 (Mészáros et al., 2009). IUPred3 is used to predict intrinsically disordered regions, while ANCHOR2 detects regions that may become structured upon interaction with binding partners. The predicted disorder profile of full-length PMI1 (Supplementary Fig. S3) is consistent with AlphaFold2 and previous reports, identifying the N-terminal and middle regions as intrinsically disordered, while the NT-C2 domain and the C-terminal DUF are predominantly structured. Notably, residues 85–119 show high disorder scores in IUPred3 but low scores in ANCHOR2, suggesting a disorder-to-order transition upon partner binding that could modulate surface charge distribution and potentially interfere with positively charged regions of the FL-NT-C2 domain, thereby regulating its membrane-binding affinity.

Although the molecular details of phosphoinositide–protein interactions remain poorly defined, structural analyses indicate that binding typically involves clusters of basic residues, often accompanied by aromatic and hydrophobic amino acids rather than single critical residues (Rosenhouse-Dantsker & Logothetis, 2007). Consistent with this model, the N- and C-terminal extensions of the FL-NT-C2 domain contain multiple lysine and arginine residues as well as aromatic and hydrophobic amino acids, and mutagenesis revealed that electrostatic interactions mediated by basic residues are essential for plasma membrane association, whereas individual aromatic substitutions had little effect on localization. While tryptophan residues did not appear to be strictly required for membrane binding, they may contribute to binding strength or domain orientation through membrane insertion or diverse noncovalent interactions. At the C-terminal extension, hydrophobic residues such as isoleucine may further stabilize membrane association, analogous to mechanisms described for other C2 domains. Together, these findings support a multistep membrane-binding model in which electrostatic interactions mediate initial membrane recruitment of the FL-NT-C2 domain, followed by stabilization through hydrophobic residue insertion, although the precise contribution and dynamics of individual residues will require further investigation.

PIPs are low-abundance signaling lipids that play central roles in membrane organization and protein targeting in plant cells (Meijer & Munnik, 2003; Noack & Jaillais, 2020). PI4P is the most abundant phosphoinositide in plants and is predominantly localized at the plasma membrane, with a smaller pool present at the trans-Golgi network and endosomal compartments; unlike in animal cells, plasma membrane-localized PI4P is the principal contributor to the negative surface charge in plants (Platre et al., 2018; Simon et al., 2016; Synek et al., 2021). This electrostatic property enables recruitment of polybasic proteins and lipid-binding domains that regulate membrane identity and trafficking. PI(4,5)P2 is also enriched at the plasma membrane, where it plays key roles in cytoskeleton-dependent processes, including cytokinesis, polarized growth, and plant–microbe interactions, and contributes to membrane curvature and vesicle formation (for review see: Heilmann & Heilmann, 2025).

We tried to identify the primary phosphoinositide responsible for PMI1 plasma membrane localization using a combination of co-localization analyses, pharmacological inhibition of phosphoinositide kinases, and targeted lipid depletion approaches. Phosphoinositide kinase inhibitors can differ in their specificity, as mammalian PI4K isoforms display variable sensitivity to wortmannin and phenylarsine oxide (PAO), with type-II PI4Ks and PI4KIIIα showing low or no sensitivity to wortmannin, whereas PI4KIIIβ is wortmannin-sensitive, and PAO preferentially inhibits type-III enzymes (Balla et al., 2002, 2005; Barylko et al., 2001; Gregory J. Downing et al., 1996; Nakanishi et al., 1995). In addition, interpretation of these experiments is complicated by the metabolic relationship between PI4P and PI(4,5)P2, as depletion of PI4P inevitably results in reduced PI(4,5)P2 levels (Simon et al., 2016). Taken together with our experimental data, these considerations suggest that PMI1 is capable of interacting with both PI4P and PI(4,5)P2, but that PI4P depletion has a more pronounced effect on PMI1 membrane association.

Given PI(4,5)P2 involvement in establishing cell polarity (Heilmann & Heilmann, 2025), it was tempting to speculate that light-dependent redistribution of phosphoinositides within the plasma membrane could underlie the observed relocalization of PMI1. However, our experimental data do not support this hypothesis, as PI4P and PI(4,5)P2 markers used in this study did not exhibit detectable relocalization within the plasma membrane upon blue-light exposure. These results indicate that bulk phosphoinositide redistribution is unlikely to be the primary driver of PMI1 displacement.

Both PI4P and PI(4,5)P2 are enriched in plasma membrane lipid nanodomains, with up to 50% of membrane PIPs localized to these structures (Furt et al., 2010). In plants, PI(4,5)P2 forms clusters with specific proteins and lipids that independently regulate distinct membrane-associated processes (Heilmann & Heilmann, 2025), and the PI4P-producing kinase PI4Kα1 localizes to nanodomains where it functions in development (Noack et al., 2022). Similarly, light activation induces clustering of the photoreceptor phot1 within plasma membrane microdomains (Xue et al., 2018). Moreover, both phototropins were shown to form homo- and heterodimers (Sztatelman et al., 2016) and phot1 transphosphorylation was reported (Kaiserli et al., 2009). Together with our results, these observations suggest that chloroplast movement signaling may be organized at plasma membrane lipid nanodomains, where PIPs could facilitate the compartmentalization of PMI1 and its association with actin regulators, photoreceptors, and other movement-related proteins.

Here, we provide evidence supporting a structural role for phosphoinositides in the regulation of chloroplast movement. Previous studies have implicated phosphoinositides in chloroplast movement primarily through pharmacological approaches, with effects largely attributed to their signaling functions via the phosphoinositide–phospholipase C (PI–PLC) pathway and downstream calcium mobilization (Aggarwal, Łabuz, et al., 2013; Anielska-Mazur et al., 2009; Grabalska & Malec, 2004). In *Lemna trisulca*, wortmannin inhibited chloroplast accumulation at low concentrations (100 nM) and avoidance at higher concentrations (1 µM), suggesting distinct regulatory mechanisms for the two responses, potentially involving PI3P in accumulation and PI4P in avoidance. In addition, PLC inhibitors blocked both accumulation and phot1-mediated avoidance responses, and weak blue light was shown to increase PI(4,5)P2 levels (Aggarwal, Łabuz, et al., 2013).

Blue light also induces cytosolic Ca^2+^ transients in *Arabidopsis* leaves in a phototropin-dependent manner (Harada et al., 2003; Harada & Shimazaki, 2009). Both phot1- and phot2-mediated Ca^2+^ increases require extracellular Ca^2+^ and are thought to involve modulation of plasma membrane Ca^2+^ channels, with PLC-dependent phosphoinositide signaling contributing specifically to phot2-induced Ca^2+^ influx. Together, these findings support a well-established role for phosphoinositides in chloroplast movement. Our results extend this model by indicating that phosphoinositides also play a direct structural role at the plasma membrane, providing a platform for the spatial organization of chloroplast movement machinery.

Cytosolic calcium levels strongly influence PMI1 localization, as demonstrated by calcium depletion and inhibitor experiments. The NT-C2 domain of PMI1 is capable of binding Ca^2+^, and calcium availability modulates its lipid-binding specificity, with preferential binding to PI4P in the presence of calcium and to PI(4,5)P2 under calcium-depleted conditions. Consistent with this, depletion of cytoplasmic Ca^2+^, as well as pharmacological inhibition of both extracellular Ca^2+^ influx and intracellular Ca^2+^ release, abolished the light-dependent redistribution of PMI1, indicating that calcium is a key regulator of this process. Although in our system fine filamentous PMI1 structures were not readily detected in darkness under standard confocal imaging conditions, likely due to single-plane acquisition, they became apparent in high-magnification z-stack images. Notably, calcium depletion resulted in the formation of thicker PMI1-associated fibers and prevented its light-induced relocalization, further supporting a role for Ca^2+^ in regulating PMI1 membrane organization and dynamics.

Functionally, the NT-C2 domain resides within the region of PMI1 that is essential for chloroplast movement, together with the N-terminal and middle unstructured regions. The C-terminal DUF appears largely dispensable for this process, although lines lacking only this domain (*pmi1* 5.12.7 and *pmi1* 5.12.12.1) retain minor defects in chloroplast movement, indicating a possible modulatory role. The functional importance of the NT-C2 domain is expected, given its role in plasma membrane localization of PMI1. Importantly, our results also highlight a critical contribution of the unstructured regions, particularly the middle segment, as plants expressing only the N-terminal unstructured region and the NT-C2 domain phenocopy the full knockout lines. This finding suggests that intrinsically disordered regions of PMI1 play an active and essential role in the regulation of chloroplast movement.

## Methods

### Plant lines and growth conditions

All *Arabidopsis thaliana* lines, including wild type (Col-0), CRISPR–Cas9 *pmi1* mutants, and transgenic reporter lines, were in the Columbia-0 background. PIP-line (Simon et al., 2014) and iDePP (Doumane et al., 2021) seeds were obtained from the Nottingham Arabidopsis Stock Centre (NASC, University of Nottingham, UK).

For chloroplast movement microscopic observations and transient expression assays, plants were grown on Jiffy peat pellets (Jiffy Products, Norway) under short-day conditions (8 h light/16 h dark, 22/19°C) in a Percival growth chamber (CLF PlantClimatics, Germany). For chloroplast movements, transmittance based measurements, under short-day conditions (10 h light/14 h dark, 22/19°C) in a Sanyo growth chamber. Seeds were stratified for 3 days at 4°C before germination.

For PAO (phenylarsine oxide) treatment, seeds were surface sterilized with chlorine gas (100 mL sodium hypochlorite + 5 mL concentrated HCl for 3 h) and aerated for 30 min. Sterilized seeds were sown on ½ MS medium (1% sucrose, 7 g/L Agargel; Sigma-Aldrich, USA), stratified for 3 days at 4°C, and grown under long-day conditions (16 h light/8 h dark, 22/19°C).

For calcium-related experiments, PMI1-YFP seeds were sterilized in 70% ethanol (5 min) followed by 50% bleach (5 min), then sown on Gamborg’s B5 medium with vitamins and 1% sucrose. After 2 days of stratification at 4°C, seedlings were grown under short-day conditions (8 h light/16 h dark, 21/17°C), and two-week-old plants were used for analyses.

### Design and Generation of *PMI1* Deletion Lines Using CRISPR/Cas9

Ten guide RNA (gRNA) sequences were designed using CRISPR-P v2.0 (Liu et al., 2017) to generate either a complete deletion of the *PMI1* gene or partial deletions of approximately 900, 1500, or 2100 nt from the 3′ end of the coding sequence. Each gRNA was 20 bp in length and included a 3′ protospacer-adjacent motif (PAM; 5′-NGG-3′) and a guanine at the 5′ end (Supplemental Table 1).

Binary vectors carrying the *Cas9* nuclease and paired gRNA sequences were constructed as follows. gRNA cassettes were amplified by PCR from plasmid pHEE2E-TRI (Wang et al., 2015; Addgene #71288) using gRNA-specific primers (Supplemental Table 1). The resulting PCR products contained both gRNA sequences, the sgRNA scaffold for the first gRNA, the U6 terminator for the first gRNA, the U6 promoter for the second gRNA, and Eco31I restriction sites. PCR products were digested with Eco31I and ligated into the pKI1.1.R vector (Tsutsui & Higashiyama, 2017; Addgene #85808) pre-digested with AarI. Correct insertion of gRNA pairs was verified by sequencing.

Validated constructs carrying *Cas9*, paired gRNAs, and the oleosin–RFP marker were introduced into *Agrobacterium tumefaciens* strain GV3101 and used to transform wild-type *Arabidopsis thaliana* (Col-0) plants via the floral dip method (Bent, 2006). Transformed seeds were selected based on red fluorescence and genotyped by PCR to assess *PMI1* deletion events. Non-fluorescent progeny were selected in subsequent generations to identify plants lacking the transgene. Stable *PMI1* edits were confirmed by PCR amplification and sequencing of the targeted genomic region.

### Generation of PMI1 Expression, Localization, and Mutant Constructs

Constructs for intracellular localization and co-localization studies were generated using the Gateway cloning system. *PMI1* cDNA fragments were cloned into entry vectors by restriction enzyme cloning (primers listed in Supplemental Table 1), verified by sequencing, and recombined into pH7CWG2, pH7WGC2, pH7YWG2 (Karimi et al., 2005), and pSITE-2CA, pSITE-2NA, pSITE-4CA, and pSITE-4NA destination vectors (Martin et al., 2009).

For recombinant protein expression, *PMI1* coding sequences were cloned into pCIOX (Addgene #51300) using overlap-assembly cloning and into pDest15 via Gateway recombination. A construct lacking the FL-NT-C2 domain was generated by PCR-based deletion from an entry clone, followed by DpnI treatment and ligation, and recombined into pSITE-2NA (Martin et al., 2009).

Constructs expressing FL-NT-C2 variants, including FL-NT-C2^WW-L^ and alanine substitutions at positions K119 and K120, were generated using the QuikChange site-directed mutagenesis method (Agilent Technologies). Additional alanine substitutions (K126A, R129A, and R133A) were introduced by a three-step overlap-extension PCR procedure (Hilgarth & Lanigan, 2020). Y294A and I293A substitutions were introduced by PCR using mutagenic primers flanking the FL-NT-C2 coding region, followed by cloning into entry vectors and Gateway recombination into destination vectors.

### Agrobacterium-mediated transient expression in *Arabidopsis thaliana* leaves

Agroinfiltration was conducted as described by Mangano et al. (2014) and (Waadt et al. (2014). *Agrobacterium tumefaciens* overnight cultures were pelleted, resuspended in induction medium (0.1% (NH4)2SO4, 0.45% KH2PO4, 1% K2HPO4, 0.05% sodium citrate, 0.2% sucrose, 0.5% glycerol, 1 mM MgSO4, pH 5.7) supplemented with antibiotics and 100 µM acetosyringone, and incubated for 5–6 h at 28 °C with shaking. Cultures were adjusted to OD600 = 0.5 (localization) or 0.4 (BiFC/co-localization), pelleted, and resuspended in infiltration buffer (10 mM MES, 10 mM MgSO4, pH 5.7) containing 200 µM acetosyringone. The suspension was infiltrated into the leaves of 4–7-week-old *Arabidopsis thaliana* plants using a needleless 2 mL syringe. Plants were maintained under standard growth conditions for 3–5 days before imaging with a confocal laser scanning microscope.

### Stress and Chemical Treatments

To assess protein localization under various treatments, transient expression in *Arabidopsis thaliana* leaves was performed as described above. For PAO treatment, PAO (P3075, Sigma-Aldrich) stock was prepared in DMSO. Leaves were infiltrated with 60 µM PAO in water for 30–40 min prior to confocal imaging. For root assays, 5–7-day-old seedlings expressing PMI1-YFP, CITRINE-2xPH(FAPP1), or CITRINE-2xPH(PLC) were incubated in 60 µM PAO solution in MS medium (1% sucrose) for 30 min with gentle rocking before imaging. Mock controls contained DMSO. For wortmannin treatment, RFP-FL-NT-C2 was transiently expressed in PIP-line plants expressing CITRINE-2xPH(FAPP1); leaves were infiltrated with 30 µM wortmannin three days post-infiltration and observed after 90 min. For dexamethasone induction, CFP-FL-NT-C2 was expressed in iDePP plants, and 24 h later, leaves were sprayed with 5 µM DEX containing 0.05% Silwet; imaging was performed after an additional 2 days.

For calcium modulation assays, leaves of two-week-old, dark-adapted (≥16 h) seedlings expressing PMI1-YFP were syringe-infiltrated with 10 mM PIPES buffer (pH 6.8) alone (control) or supplemented with one of the following treatments: 10 µM A23187 + 1 mM EGTA or 10 µM A23187 + 10 mM Ca²⁺ for 1–1.5 h, 20 µM verapamil for 2 h, or 15 µM thapsigargin for 2–30 min. Treatment conditions and incubation times were based on established protocols (Anielska-Mazur et al., 2009; Tlałka & Fricker, 1999).

Samples were prepared under physiological darkness, infiltrated with the respective solutions, and maintained in the same medium until imaging. Confocal images were acquired using 514 nm excitation, and blue light treatment was applied with a 458 nm laser for 2.5 min.

### Confocal microscopy

Most imaging was performed using a Nikon C1 laser-scanning confocal system on a TE2000E platform (Nikon Instruments, Amsterdam, The Netherlands). Excitation/emission settings were as follows: CFP, 409 nm/450–485 nm; GFP/YFP, 488 nm/535 ±15 nm; RFP, 543 nm/605 ±37 nm; chlorophyll autofluorescence, 638 nm/≥650 nm. A 60× Plan-Apochromat oil immersion objective was used, except for chloroplast movement assays, where a 40× Plan Fluor oil immersion objective was applied.

Observations of calcium-modulating treatments were conducted using an Axio Observer.Z1 inverted microscope (Carl Zeiss, Jena, Germany) equipped with an LSM 880 confocal module. Blue laser light (458 nm), emitted by an argon-ion laser was applied for stimulation, and YFP fluorescence was recorded at 514 nm excitation with emission detection from 519–620 nm. Chlorophyll autofluorescence was excited at 633 nm and detected at 647–721 nm using a long-distance 25× Plan-Apochromat (NA 0.8) multi-immersion objective with glycerol immersion.

### Protein Expression and Purification

GST-tagged core-NT-C2, ΔN-NT-C2, ΔC-NT-C2, FL-NT-C2, and GST alone were expressed in *E. coli* Rosetta (DE3) pLysS carrying pDEST15 constructs containing the respective coding sequences or a STOP codon (for GST). Expression was induced with 0.5 mM IPTG for 2 h at 37 °C. Clarified lysates were purified using an FPLC system with an EconoFit Profinity GST column (Bio-Rad, USA), washed with TBS (pH 7.6), and eluted with TBS containing 20 mM reduced glutathione (Carl Roth, Germany). Protein purity was verified by 4–20% SDS–PAGE (Supplementary Figure S4), and samples were concentrated and stored at −80 °C.

### Protein-Lipid Overlay

Membrane Lipid Strips (P-6002), PIP Strips (P-6001), and PIP Arrays (P-6100) (Echelon Biosciences, USA) and GST-tagged recombinant proteins were used following the protocol of Shirey et al. (2017). Before blocking, 0.05 and/or 0.5 pmol of the tested protein and GST alone were spotted on the membrane as a detection control. Membranes were blocked in TBS-T containing 3% BSA (Sigma-Aldrich, USA) for 1 h at room temperature, then incubated overnight at 4 °C with 0.5 μg/mL protein in the blocking buffer. After washing with TBS-T, bound proteins were detected using anti-GST antibody (1:500; Sigma-Aldrich, Cat. No. G1160) for 2 h, followed by HRP-conjugated anti-mouse secondary antibody (1:25,000; Invitrogen, Cat. No. 31457) for 1 h. Signals were visualized by chemiluminescence using the Pierce ECL substrate (Thermo Scientific, USA). The PLO assay with EGTA was performed as described above, except that blocking, protein, and wash buffers were supplemented with 5 mM EGTA.

### Calcium Binding Assay using Fura-2

GST-tagged FL-NT-C2 and GST (control) were clarified by centrifugation and processed in Protein LoBind tubes (Eppendorf, Germany). Proteins were treated with 5 mM EGTA for 1 h to remove residual calcium and desalted using Zeba Spin Desalting Columns (7K MWCO, Thermo Scientific, USA). The assay was performed as described by Jing et al., (2016) with minor modifications. Glutathione Sepharose 4B (GE Healthcare/Cytiva, USA) was equilibrated, and 100 µL of 50% slurry was mixed with 50 µL of calcium buffer (TBS + 0.2 mM CaCl2) and 150 µL of protein solution (0.67–2.6 µM final concentration; 33 µM Ca^2+^ total). Samples were incubated for 30 min at room temperature with gentle rotation, centrifuged (500 × g, 5 min), and 160 µL of supernatant was collected. Free Ca^2+^ was quantified using Fura-2 (Cayman Chemical, USA): 75 µL of supernatant was mixed with 75 µL TBS and 5 µL of 1 mM Fura-2, incubated 5 min in the dark, and fluorescence was recorded (excitation 350 nm, emission 506 nm; SpectraMax iD3, Molecular Devices, USA). Data from five independent experiments were analyzed by Student’s *t*-test in JASP (v0.19.0.0, University of Amsterdam); Bonferroni correction for four comparisons was used. Graphs were prepared using Microsoft Excel 2019.

### Microscopic Analysis of Chloroplast Positioning

Adult mutant plants were dark-adapted for at least 16 h, and all sample preparations were conducted under dim green light. Detached leaves were syringe-infiltrated with 10 mM PIPES buffer (pH 6.8) to reduce light scattering and improve optical clarity. Control samples were kept in darkness until imaging, while others were illuminated with either high-irradiance (100 μmol m^−2^ s^-1^) or low-irradiance (2.3 μmol m^−2^ s^-1^) blue light for 1 h before observation. Optical cross-sections of the palisade mesophyll were acquired, and Z-stack projections were generated using ImageJ software (Schneider et al., 2012).

### Quantitative Analysis of Chloroplast Movements Using Light Transmittance

Leaf transmittance changes were examined by the photometric method (Gabryś et al., 2017), with a custom photometric setup. Chloroplast movements were triggered with blue light of 1 or 100 µmol m^−2^ s^-1^ (peak at 455 nm, M455L4 LED, Thorlabs).

The red measuring light (peak at 660 nm, M660L4 LED, Thorlabs) was modulated at 1033 Hz. The light beams were combined with a dichroic mirror (DMLP550, Thorlabs) and directed towards the detached leaf, mounted in front of one of the ports of an integrating sphere (IS200-4, Thorlabs). At another port of the sphere, a photodiode detector (DET100A2, Thorlabs) was mounted. Leaf transmittance curves were analyzed with a custom script by Mathematica (Wolfram Research). Irradiance values were measured with the LI-190R sensor (LI-COR) and the Keithley 6485 picoampere meter. 10-12 leaves were measured for each group in seven (1 µmol m^−2^ s^-1^) or four (100 µmol m^−2^ s^-1^) batches. The statistical analysis was performed in the R Software (R Core Team, 2025). The amplitude of transmittance changes, the rate of transmittance changes and the transmittance of a leaf in the dark-adapted state were treated as response variables. A mixed linear model was fitted with the *nlme* package (Pinheiro et al., 2023; Pinheiro & Bates, 2000), explaining the relation between a response and the plant line, treated as a categorical fixed factor. In addition, the model contained a plant batch as a random factor. The model accounted for the possibility of unequal variances between different plant lines. The differences in mean values of the responses between plant lines were examined for statistical significance with Tukey’s method, using a combination of the *emmeans* (Lenth, 2025) and *multcomp* packages (Hothorn et al., 2008).

## Author contributions

O.S., D.C., and G.D. conceived the work and designed the experiments. D.C performed most of the experiments described in the study. O.S. prepared PMI1-YFP expressing plants and performed pilot experiments. Z.S. carried out the construction of CRISPR Cas9 mutants and performed their initial characterization. J. Ł. and P. H. performed chloroplast movement analysis using red light transmittance measurements. P.H. and D.C performed the microscopic experiments analyzing calcium impact on PMI1 displacement. O.S and D.C. obtained the funding. O.S and D.C. wrote the original text draft, and all authors contributed to the manuscript and discussed the results.

## Acknowledgements

We would like to thank Dr Lien Brzeźniak (IBB PAS) for help with genome editing and Adrian Kasztelan (IBB PAS) for help with setting up protein purification. The authors used ChatGPT (OpenAI) to assist with language editing and improvement of the manuscript. The authors take full responsibility for the content of the publication

