## Supplementary material for "NT-C2–Dependent Phosphoinositide Binding Controls PLASTID MOVEMENT IMPAIRED1 Localization and Function": Suplementary materials

**Supplementary table 1 Primers used in the study**

| Name | Sequence (5' → 3') |
|---|---|
| Cloning of PMI1 and its domains | |
| PMI1-N-unstr:C-NotI-F | TCTGCGGCCGCGATGGCAGGAGAATATTCCGG |
| PMI1-N-unstr:C-KpnI-R | TCTGGTACCAACCAGCCCTCGAATCGGC |
| PMI1-N-unstr:N-NotI-F | TCTGCGGCCGCGGCAGGAGAATATTCCGGTAGC |
| PMI1-N-unstr:N-KpnI-R | TCTGGTACCCTAAACCAGCCCTCGAATCGGC |
| PMI1-NTC2:C-NotI-F | TCTGCGGCCGCGATGCGGATTGGTATGCAGAAGCTAAGC |
| PMI1-NTC2:C-KpnI-R | TCTGGTACCGTCTTTCTCCATAATCTGAAACCC |
| PMI1-NTC2:N-NotI-F | TCTGCGGCCGCGCGGATTGGTATGCAGAAGCTAAGC |
| PMI1-NTC2:N-KpnI-R | TCTGGTACCCTAGTCTTTCTCCATAATCTGAAACCC |
| PMI1-M-unstr:C-NotI-F | TCTGCGGCCGCGATGGGTGGAGCTGGGATTTACAG |
| PMI1-M-unstr:C-KpnI-R | TCTGGTACCCGAGAGGTAATTCTCCGATTC |
| PMI1-M-unstr:N-NotI-F | TCTGCGGCCGCGGGTGGAGCTGGGATTTACAG |
| PMI1-M-unstr:N-KpnI-R | TCTGGTACCCTACGAGAGGTAATTCTCCGATTC |
| PMI1-CDUF:C-EcoRI-F | ATGAATTCATGTCGGATCTCGGTAAAGGCATTG |
| PMI1-CDUF:C-SalI-R | TACTGTCGACATGCAATTTCACATCAGGGTTC |
| PMI1-extended-NTC2:C-F | TCTGCGGCCGCGATGAAGAAAGGGATTTGGAATTGGAAG |
| PMI1-extended-NTC2:C-R | TCTGGTACCACTGTAAATCCCAGCTCCACC |
| PMI1-extended-NTC2:N-F | TCTGCGGCCGCGAAGAAAGGGATTTGGAATTGGAAG |
| PMI1-extended-NTC2:N-R | TCTGGTACCCTAACTGTAAATCCCAGCTCCACC |
| PMI1noC2-rev | CTCTTCCTTCACACCCGAACCCGATGACG |
| PMI1noC2-fwd | AGTAAACAAGGCGAATTCGGGATGAAACCG |
| pCIOX-lin-rev | ACCACCAATCTGTTCTCTGTGAG |
| pCIOX-lin-for | CGCGGATCCGAATTCGAG |

| | |
|---|---|
| C2-pCIOX-SLIC-for | CACAGAGAACAGATTGGTGGTAAGAAAGGGATTTGGAATTGGAAG |
| C2-pCIOX-SLIC-rev | GAGCTCGAATTCGGATCCGCGCTAACTGTAAATCCCAGCTCCACCG |
| Site directed mutagenesis of NT-C2 domain | |
| K126A-R129A-R133A-antisense | CTGCATACCAATCGCAACCAGCCCTGCAATCGGCGCCCAATTCCAAATCCC |
| K119A-K120A | CAGGCTCCGCGGCCGCGGCGGCAGGGATTTGGAATTGG |
| K119A-K120A-antisense | CCAATTCCAAATCCCTGCCGCCGCGGCCGCGGAGCCTG |
| K126A-R129A-R133A | GGGATTTGGAATTGGGCGCCGATTGCAGGGCTGGTTGCGATTGGTATGCAG |
| W123L-W125L-antisense | CGAATCGGCTTCAAATTCAAAATCCCTTTCTTCGCGGCC |
| W123L-W125L | GGCCGCGAAGAAAGGGATTTTGAATTTGAAGCCGATTCG |
| Y294A-antisense | CCGGTACCCTAACTGGCAATCCCAGCTCCACCG |
| Y294A | CGGTGGAGCTGGGATTGCCAGTTAGGGTACCGG |
| Cloning of gRNA for CRISPR/Cas9 genome editing | |
| PMI1g30F | GGTCTCGATTGTAGGAAGTTTGCACATTGGgttttagagctagaaatagcaag |
| PMI1g36F | GGTCTCGATTGGAGGCTTACCGCGGCGCAAgttttagagctagaaatagcaag |
| PMI1g66R | GGTCTCGAAACCAATCTGTTTGGAGATTTGcaatctcttagtcgactctacc |
| PMI1g207R | GGTCTCGAAACCACAAACCCCAAATAAACCcaatctcttagtcgactctacc |
| PMI1g153R | GGTCTCGAAACGACCGCAGTTCAGAGATGTcaatctcttagtcgactctacc |
| PMI1g218R | GGTCTCGAAACCTCGCTGGCGTCCACGCCTcaatctcttagtcgactctacc |
| PMI1g275R | GGTCTCGAAACTCACACCCGAACCCGATGAcaatctcttagtcgactctacc |

| | |
|---|---|
| PMI1g84R | GGTCTCGAAACCGACATTCTAAATCCTGATcaatctcttagtcgactctacc |
| Identification of *pmi1* mutants | |
| PMI1_For1 | ACCCTCAGCTTCCTTATCCC |
| PMI1_For2 | GAGTTGAGAGAAGGCCACG |
| PMI1_For3 | CTGATCCAATCGCTTCACC |
| PMI1_For4 | GGATTTGGAATTGGAAGCCG |
| PMI1_For5 | ATGAAGTGAGTACAGCACGA |
| PMI1_For6 | TGAGTTGATTCAGGAGTCGG |
| PMI1_For7 | GCATGTGGTGGATTTGAGTG |
| PMI1_For8 | GGAGGTGATGGTGAAACAG |
| PMI1_For9 | GGATGAAGAAACCGAGAAGC |
| PMI1_Rev1 | AACATTAGGGAAAATTAGTTGAACC |
| PMI1_Rev2 | GCAATTTCACATCAGGGTTCC |
| PMI1_Rev3 | CAGCACCGTCTTTAGTTTCC |
| PMI1_Rev4 | AACCCTGAGAAACCCTACAC |
| PMI1_Rev5 | AAACTGACTCCACACCACTC |
| PMI1_Rev6 | GATTCAAATGCTCCATCCCG |
| PMI1_Rev7 | AATTCAGGCCTTACCACAATC |
| PMI1_Rev8 | TTCAGATTCCTGTTGTTGTAGG |

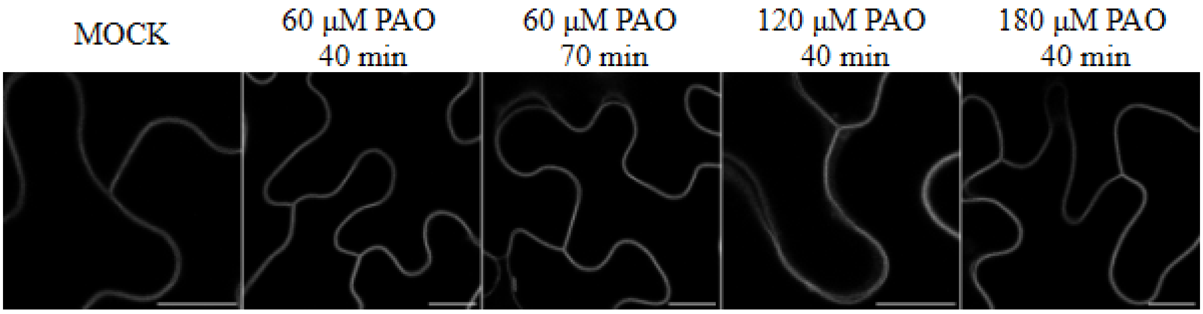


**Supplementary Figure S1. Influence of PAO on the localization of PIP4 markers in adult leaf epidermis**

The PI4P marker CITRINE–2xPH(FAPP1) shows no detectable change in localization in leaf epidermal cells following treatment with 60 µM phenylarsine oxide (PAO), a PI4K inhibitor, even after prolonged incubation or increased inhibitor concentration. Scale bar = 10 µm

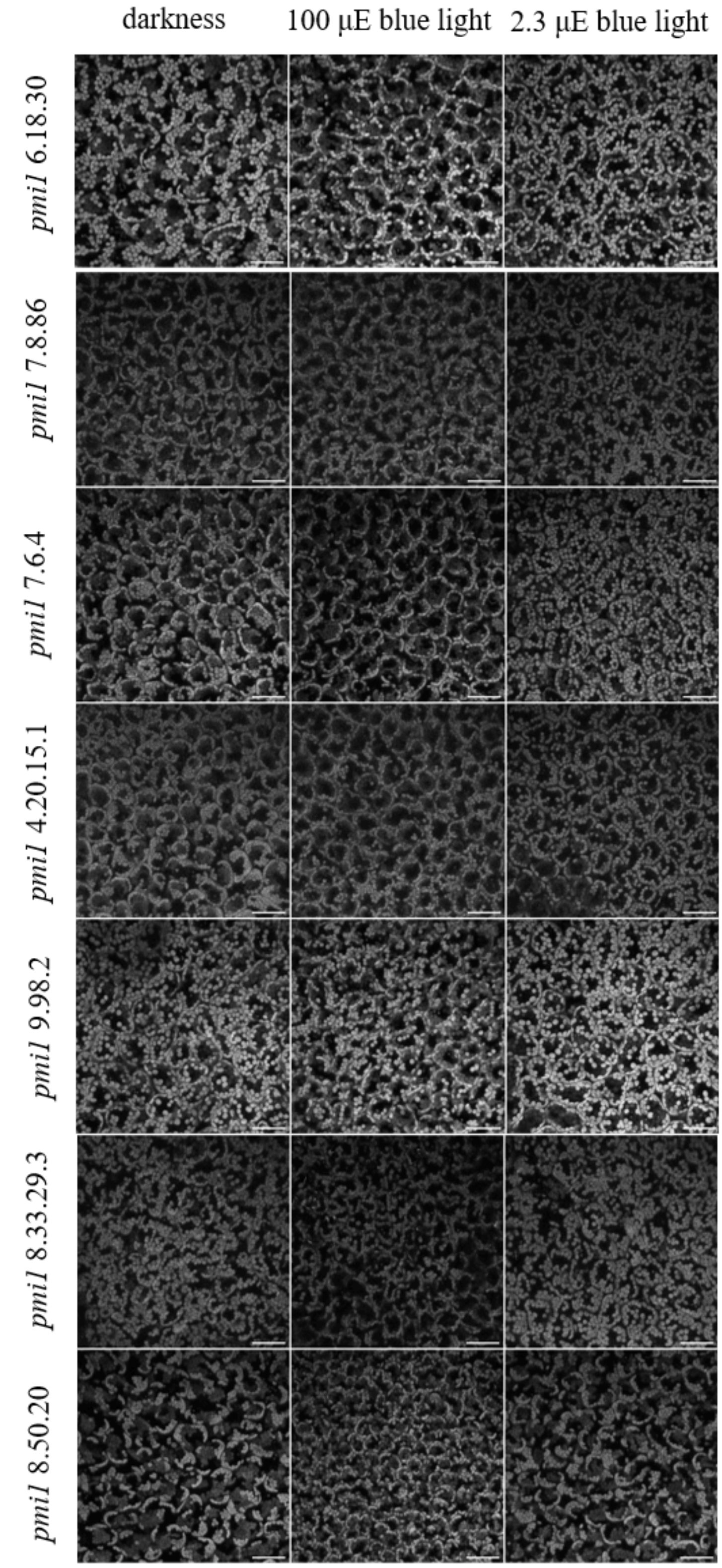


**Supplementary Figure S2. Chloroplast movements in *pmi1* mutant plants visualized by confocal microscopy using chlorophyll autofluorescence**

Mutants not included in Figure 6 are presented. Scale bar = 50 µm. Images shown are representative of ten independent biological replicates.

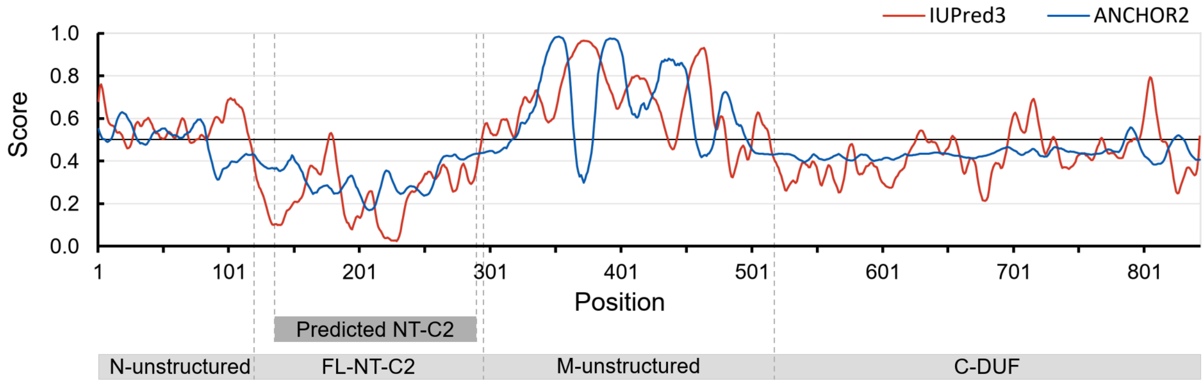


**Supplementary Figure S3. Predictions of disordered regions in PMI1 protein**

IUPred3 and ANCHOR2 predictions of disordered regions of full-length PMI1 in unbound and bound form, respectively. A score of 0.5 denotes the threshold between ordered and disordered regions.

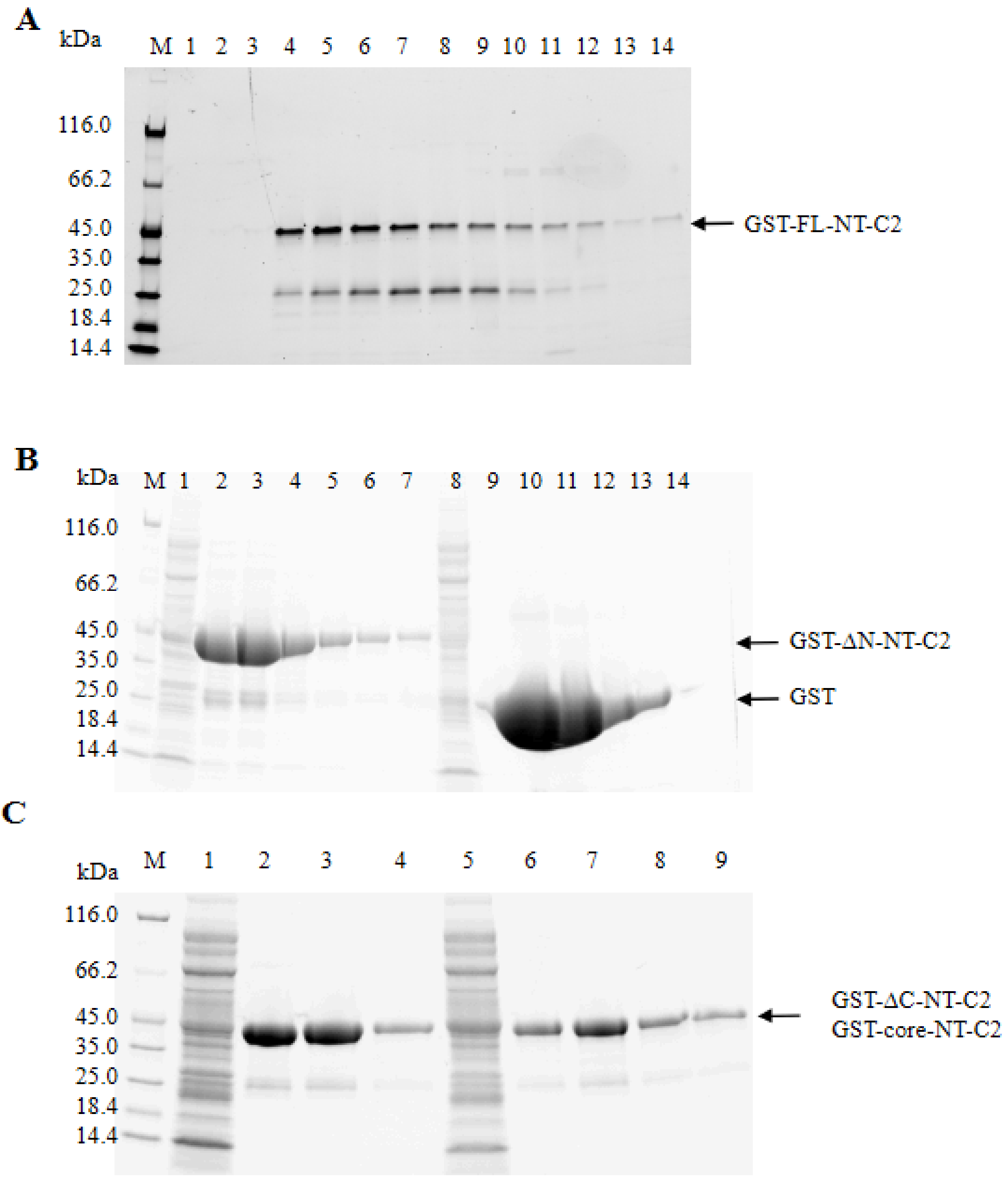


**Supplementary Figure S4. Purification results of GST-tagged FL-NT-C2, core-NT-C2, ΔN-NT-C2, ΔC-NT-C2, and GST**

4–20% SDS-PAGE analysis of (A) fractions after ion exchange of the GST-FL-NT-C2 and (B–C) fractions after affinity purification of the GST-ΔN-NT-C2, GST (B), GST-core NT-C2, and GST-ΔC-NT-C2(C), respectively. The GST-tagged proteins were all expressed from Rosetta DE3 pLys *E. coli* strain bearing pDest15 plasmid with appropriate coding sequence or STOP codon (for GST). Bacteria were grown in 500 mL TB medium supplemented with ampicillin (100 µg/mL) and chloramphenicol (25 µg/mL) at 37 °C to an $OD_{600}$ of 0.7 and protein expression was induced with 0.5 mM IPTG for 2 h. Cells were harvested, processed immediately or stored at −80 °C, and lysed in TBS (pH 7.6) containing protease inhibitors,

PMSF, DTT, lysozyme, Triton X-100, $MgCl_2$, ATP, and viscolase, followed by sonication and clarification by centrifugation. The filtered supernatant was purified by FPLC using a Profinity GST column (Bio-Rad), washed with TBS, and eluted with TBS supplemented with 20 mM reduced glutathione.  Lanes 1 and 8 in panel B and lanes 1 and 5 in panel C represent sample application fractions, and the remaining lanes correspond to eluted fractions. For GST-FL-NT-C2 additional ion exchange step was employed with EconoFit UNOsphere Q Column (Bio-Rad, Hercules, USA). Fractions presented in (A) lanes 4–6 (B) lanes 2-4 and 10-12 (C) lanes 2-4 and 6-8 were concentrated using Pierce™ Protein Concentrators PES, 30K MWCO (Thermo-Scientific, USA) and the buffer was exchanged to TBS, pH 8.0. The protein aliquots were snap-frozen and kept in -80℃.
